# Optimization of Spectral Library Size Improves DIA-MS Proteome Coverage

**DOI:** 10.1101/2020.11.24.395426

**Authors:** Weigang Ge, Xiao Liang, Fangfei Zhang, Luang Xu, Nan Xiang, Rui Sun, Wei Liu, Zhangzhi Xue, Xiao Yi, Bo Wang, Jiang Zhu, Cong Lu, Xiaolu Zhan, Lirong Chen, Yan Wu, Zhiguo Zheng, Wangang Gong, Qijun Wu, Jiekai Yu, Zhaoming Ye, Xiaodong Teng, Shiang Huang, Shu Zheng, Tong Liu, Chunhui Yuan, Tiannan Guo

## Abstract

Efficient peptide and protein identification from data-independent acquisition mass spectrometric (DIA-MS) data typically rely on an experiment-specific spectral library with a suitable size. Here, we report a computational strategy for optimizing the spectral library for a specific DIA dataset based on a comprehensive spectral library, which is accomplished by *a priori* analysis of the DIA dataset. This strategy achieved up to 44.7% increase in peptide identification and 38.1% increase in protein identification in the test dataset of six colorectal tumor samples compared with the comprehensive pan-human library strategy. We further applied this strategy to 389 carcinoma samples from 15 tumor datasets and observed up to 39.2% increase in peptide identification and 19.0% increase in protein identification. In summary, we present a computational strategy for spectral library size optimization to achieve deeper proteome coverage of DIA-MS data.

## Introduction

Data-independent acquisition mass spectrometry (DIA-MS) based proteomics coupled with targeted data analysis is playing an increasing role in biomedical studies (1), owing to its high degree of reproducibility, quantitative accuracy, and high throughput (2, 3). Both spectral library-free and library-based strategies are being applied to analyze DIA-MS data (4). While the library-free strategies (5, 6) could identify peptides directly from DIA-MS itself without the requirement of an external spectral library, the depth of proteomic coverage is limited at the moment (7–9). The more widely adopted strategy is based on building a spectral library using the corresponding data-dependent acquisition mass spectrometry (DDA-MS) datasets of the samples of interest (10), or a pre-built library from public data repositories (11–14).

The size of the spectral library has a direct impact on the performance of DIA-MS data analysis (15). A larger number of DDA-MS runs, particularly from fractionated samples, leads to a more comprehensive spectral library enabling potential detection of a larger number of peptides and proteins from the DIA-MS datasets (15). However, it also generates a larger search space and reduces the statistical power to detect true positives (16, 17). Extra concerns are raised where the proteins and peptides within the library may not be specific to a particular specimen, potentially introducing more false positives (18). Other drawbacks include the prolonged computational time which is approximately linearly correlated with the size of the library (19), and distortion of retention time (RT) distribution for alignment (20).

The spectral library size could be optimized to improve DIA-MS performance. The Van Eyk group have reported that applying a comprehensive fractionated library led to higher number of protein/peptide identifications from DIA-MS datasets than un-fractionated libraries with limited sizes (15). Similar results have been reported by Uszkoreit group, where they found larger library led to higher peptide and protein identification but the increase was minimal when the library is comprehensive enough (21). The combination of an in-house built library with external libraries from public data improves DIA data analysis performance (17). Inclusion of internal library extracted from DIA files also improved peptide and protein identification (9). On the other hand, it has also been observed that libraries of very large size led to higher FDRs in the DIA-MS analyses and hence compromises the identification results (17). It was further demonstrated that, even within the same spectral library, controlling the confidence of peptide identifications to exclude redundant peptides could improve peptide and protein identification results (16). Although these studies have repetively reported the importance of the size of spectral library size, a systematic evaluation and optimization of library size is still lacking.

Here, we propose a two-step strategy called subLib to generate the experiment-specific subset libraries using *a priori* analysis of the DIA data to improve the proteomic coverage. The strategy to derive a subset library of optimal size was further applied to analyze the DIA data of 15 human tumors.

## Materials and Methods

### Colorectal cancer dataset

To evaluate our strategy, the DIA-MS datasets were collected from a colorectal cancer proteomic project in our group (Xiang *et al.,* manuscript in preparation). Briefly, 286 FFPE samples from 44 colorectal cancer patients were processed into peptides with a pressure cycling technology (PCT)-based protocol as described in the previous study (22). They were subjected to data acquisition on the nanoflow EASY-nLC™ 1200 System coupled with Q Exactive HF hybrid Quadrupole-Orbitrap in DIA mode over a gradient of 60 min using 24 DIA windows spanning from 400 Da to 1200 Da.

### Fifteen datasets of multiple tumor types

A total of 389 tumor tissue samples from 15 tumor types were collected. The gastric carcinoma (n=30) and thyroid carcinoma (n=30) samples were collected from the First Affiliated Hospital College of Medicine, Zhejiang University. The prostate carcinoma (n=30) and bone carcinoma (n=30) samples were collected from the Second Affiliated Hospital College of Medicine, Zhejiang University. The liver carcinoma (n=33) and leukemia (n=27) samples were collected from Wuhan Union Hospital. The ovarian carcinoma (n=30) samples were collected from Zhejiang Cancer Hospital. The cervical carcinoma (n=28) samples were collected from Shengjing Hospital of China Medical University. The lung adenocarcinoma (n=32), gallbladder carcinoma (n=20), pancreatic adenocarcinoma (n=20), myosarcoma (n=19), clear cell renal cell carcinoma (CCRCC, n=20), diffuse large B-cell lymphoma (DLBCL, n=19), and papillary thyroid cancer (PTC, n=21) were collected from Harbin Medical University Cancer Hospital. All samples were approved by the ethics committees of their respective hospitals. The tissue samples were prepared with PCT-based tissue lysis and protein digestion protocol (22) and analyzed by DIA-MS, as listed in Table S1. Ethics approvals for this study were obtained from the Ethics Committee or Institutional Review.

### Proteomic data analysis workflow

The raw DIA-MS data files were converted to mzXML format using the msConvert tool in ProteomeWizard (23). The DIA-MS datasets were analyzed using the open-source software OpenSWATH (version 2.4.0) (24) with the following criteria: common internal reference peptides (CiRTs) of each tissue were applied respectively for retention time alignment; m/z extraction window was set to 30 ppm, and RT extraction window was set between 200-800 seconds, depending on different gradients of the DIA-MS module (Table S1). PyProphet (version 2.1.3) (24) was used for statistical validation via setting the global cutoff of FDR as 0.01 at both peptide and protein levels. Protein inference was performed as described previously (25). Unless otherwise mentioned, the software parameters were kept the same for all the analyses in this study.

### Subset library generation

We proposed a two-step strategy to take a subset of the spectral library. Firstly, the public library is taken to analyze the candidate DIA-MS dataset using the OpenSWATH workflow. Different FDR cutoffs were set to generate a list of identification results. Afterwards, they were matched against the public library to generate experiment-specific subset libraries.

In this study, we set the DIA Pan-Human Library (DPHL) (12) as the baseline library to analyze the colorectal cancer dataset containing 284 DIA-MS data files. FDR cutoffs were set at 0.01, 0.02, 0.03, 0.04, 0.05, 0.06, 0.07, 0.08, 0.09, 0.1, 0.2, 0.3, 0.4, 0.5 and 0.6 (n=15), to generate 15 identification results. After matching with DPHL, OpenSwathDecoyGenerator.exe in OpenMS (version2.4.0) was applied to generate equal amount of decoys in mutated fashion. The resultant subset library is a combination of DPHL subsets and decoys.

## Results and Discussions

### Generation of the subset library by refining DPHL

For data comprehensiveness and accessibility, DPHL built from 16 human tissue types containing 359,627 peptide precursors and 14,782 protein groups was used as the baseline spectral library. A DIA-MS dataset of 286 colorectal cancer sample cohort was analyzed to derive the initial identifications. We set the FDR cut-off for peptide precursor and protein identification to 0.01, 0.02, 0.03, 0.04, 0.05, 0.06, 0.07, 0.08, 0.09, 0.1, 0.2, 0.3, 0.4, 0.5 and 0.6 (a total of 15 tests), then retrieved the resultant subset libraries at each FDR cutoff. The four representative DIA-MS data files (sample A1-A4) within the cohort and two external colorectal cancer DIA-MS data files (sample B1 and B2) were taken to evaluate the identification performance of each subset library (Figure 1A). The number of identified peptides shows a generally decreasing trend as the FDR cutoff increases (Figure 1C), with the exceptions when FDR increases from 0.01 to 0,02, and from 0.04 to 0.05. The number of identified proteins increased as the FDR cut-off increased from 0.01 to 0.05, and gradually decreased afterward, with a drastic decline when the cutoff was beyond 0.1 (Figure 1D). This is not unexpected since the peptides identified with high FDR are more likely absent in the sample at the detection limit. As the library size increased, the negative effect prevailed. The best result was obtained from the library with a FDR cutoff of 0.05. The optimal library was composed of 85,655 peptide precursors, 62,390 peptides, and 6,448 protein groups, leading to the identification of 29,979 peptide precursors and 4,418 protein groups, respectively. This optimized library led to 44.7% and 38.1% increase of peptide precursors and protein groups, respectively, compared with the results by the unfiltered DPHL (Figure S1). The subset library with the FDR cutoff of 0.05 was the best subset library which was hence adopted for further evaluation. The DIA files used for library size optimization from samples A1-A4 led to similar data to those from independent samples (B1 and B2), suggesting that the library size optimization is generic and applicable to DIA files of the same tissue type.

**Figure 1.**
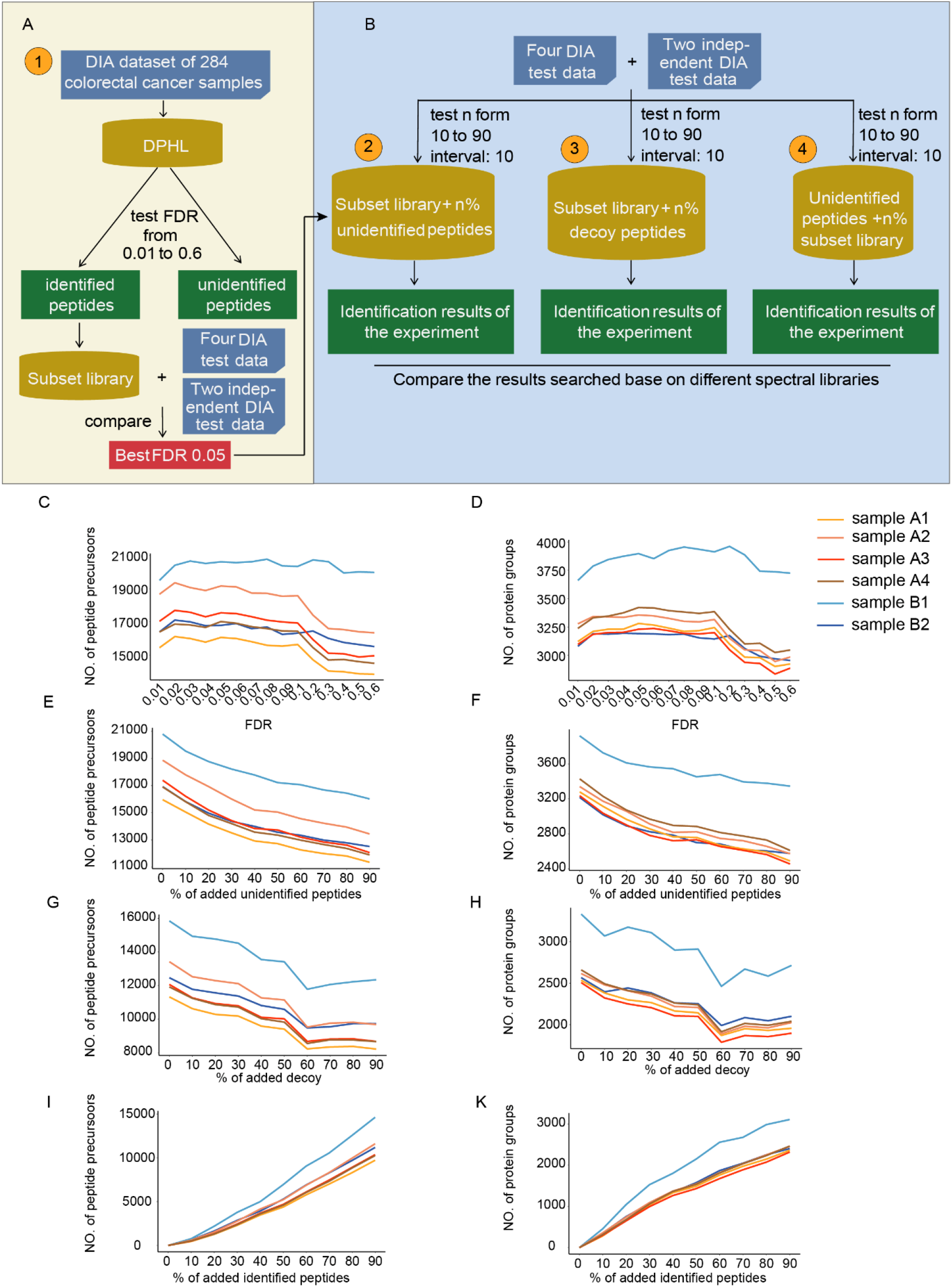
Optimizing DPHL in the DIA dataset of colorectal cancer. (A) The workflow of spectral library optimization. Step 1: Select the best FDR for refining the subset library from the public DPHL library. The subset library refined from DPHL with FDR of 0.05 is considered as the optimal subset library, which was used as a primary optimized subset spectral library in this study. Step 2: Evaluate the performance of the spectral library consisting of the subset library and n% unidentified peptides. Step 3: Evaluate the performance of spectral library consisting of subset library and n% decoy peptides. Step 4: Evaluate the performance of spectral library consisting of unidentified peptides and n% peptides from the subset library. By comparing all the identification results, the subset library refined from DPHL with FDR of 0.05 is the best experiment-specific spectral library for DIA data analysis. The numbers of identified peptides (C) and proteins (D) based on the subset libraries which were refined from DPHL at nine different FDRs. The numbers of identified peptides (E) and proteins (F) based on the spectral libraries consisting of subset library and n% unidentified peptides. The numbers of identified peptides (G) and proteins (H) based on the spectral libraries consisting of the subset library and n% decoy peptides. The numbers of identified peptides (I) and proteins (K) based on the spectral libraries consisting of unidentified peptides and n% peptides from the subset library.

### Adding unidentified peptide procursors to the subset library sacrificed identification

To check if unidentified peptide precursors in a spectral library would affect the DIA-MS proteome coverage, we randomly generated nine sets of DPHL peptides that were excluded from the subset library (defined as “unidentified peptides”), with precursor number equivalent to n% of the subset library (n=10, 20, …, 90), and combined them with the subset library peptides (Figure 1B). When applying the reconstructed spectral libraries to analyze the test DIA dataset, a steady decrease of identified peptides and proteins was observed as more unidentified precursors were included (Figure 1E, F), with the highest proteome coverage coming from the library with no unidentified peptides, summing up to 29,712 peptide precursors and 4,433 protein groups.

We also replaced the unidentified peptides to *in silico* generated decoy peptides and repeated the above analyses. Peptide/protein identifications decreased as the computational peptide proportions increase from 0% to 60%. Further addition of decoys would, however, subtly increase protein identifications (Figure 1G, H). The highest proteome coverage came from the library with no decoy interferences, summing up to 19,322 peptide precursors and 3,461 protein groups. We hence concluded that any false positive interference in the library would suppress the peptide/protein identification.

### Adding subset library peptides to interferences improves identification

We then conducted a backward analysis by adding increasing proportions of subset library peptides to the unidentified peptides (Figure 1B). The spectral library composed by precursors of unidentified peptides solely (n=0) could not identify any peptide or protein in the DIA-MS data. The numbers of identification of peptides and proteins exhibited almost marked increase as n increased (Figure 1I, K). Together with the above results, they validated the effectiveness of setting FDR cutoff as 0.05 to eliminate false positive targets.

### Applying subLib to DIA-MS of 15 tumor sample types

We named the library generation strategy “subLib” and further applied it to the fifteen DIA datasets of different types of cancer samples, including bone, cervical, DLBCL, gallbladder, gastric, leukemia, liver, lung, myosarcoma, ovarian, pancreatic, prostate, PTC, and CCRCC (Figure 2A). Peptide/protein identifications using the subset library exceeded that from using DPHL in most cases (Figure 3A), and over 99% of the protein identifications were overlapped in every cancer type (Figure S2). We collectively found that the subLib strategy outperformed the DPHL strategy in all cancer types, with the most prominent increase from PTC carcinoma samples (19.02% increase in protein groups and 36.17% increase in peptide precursors, Figure 2B). Of note, the discrimination ability to separate the targets from decoys led to a marked increase (Figure 2C), further validating that the subLib strategy can reduce false positives in clinical proteomic data. Missing values were equivalent between DPHL and the subLib strategy (Figure 3B), and the protein quantification results were in good accordance as well with Pearson correlation ratios all over 0.92 across all the tumor tissue types (Figure 3B), suggesting that decreasing library sizes by adjusting FDR values does not impair protein identification nor quantification. Moreover, different tumor types could be well resolved using the thus generated protein matrix (Figure 2D). These results indicate that this subLib strategy could be generically used for DIA data generated from different samples.

**Figure 2.**
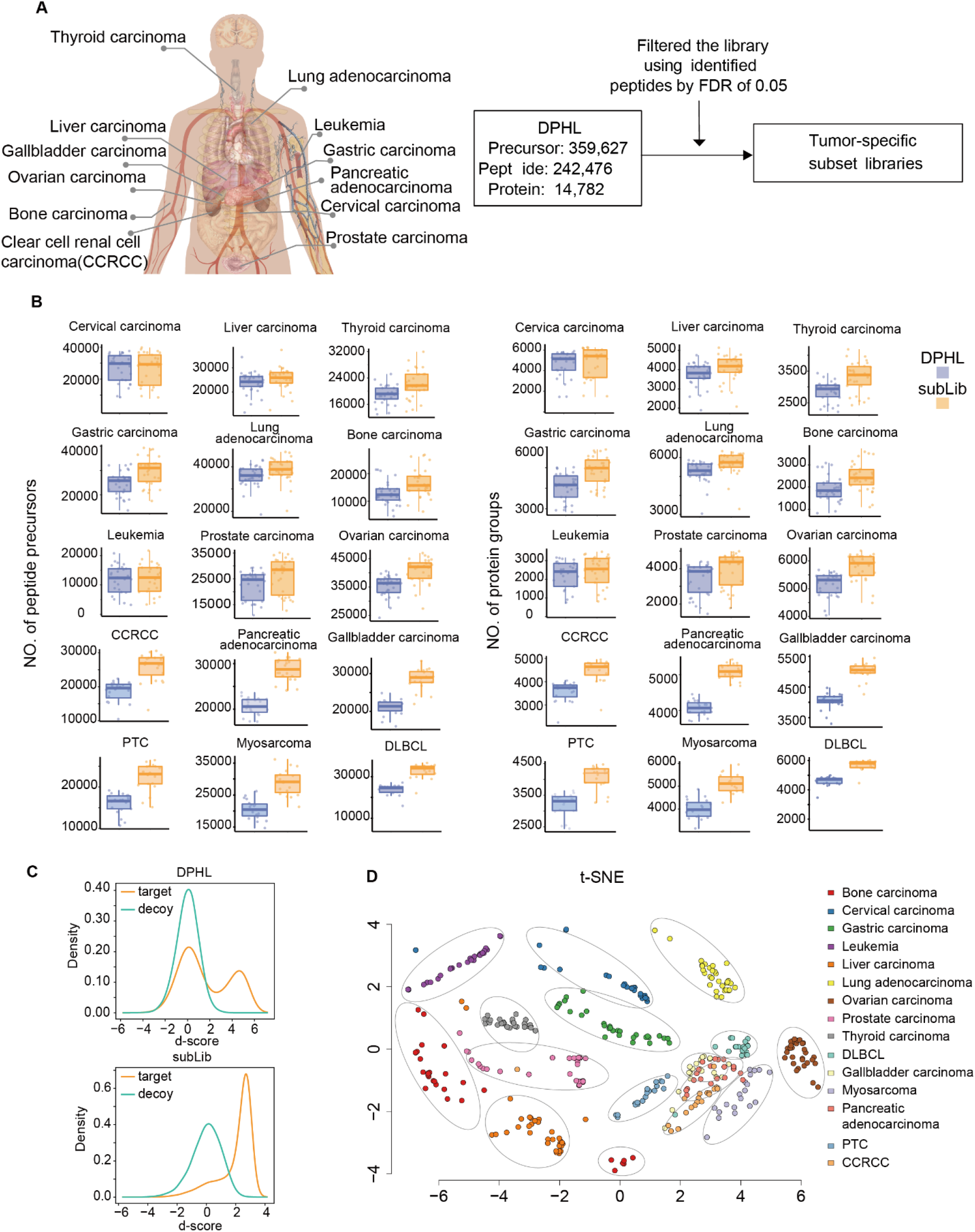
Tumor-specific subset library improves the identifications compared with DPHL. (A) The workflow of the subLib strategy. (B) The number of peptides and proteins identified base on tumor-specific subLib and DPHL in 15 tumor types. (C) The distribution of discrimination score (d-score) of the target and decoy of the subset library and DPHL. (D) The tSNE plot shows the samples are well resolved by tissue type.

**Figure 3.**
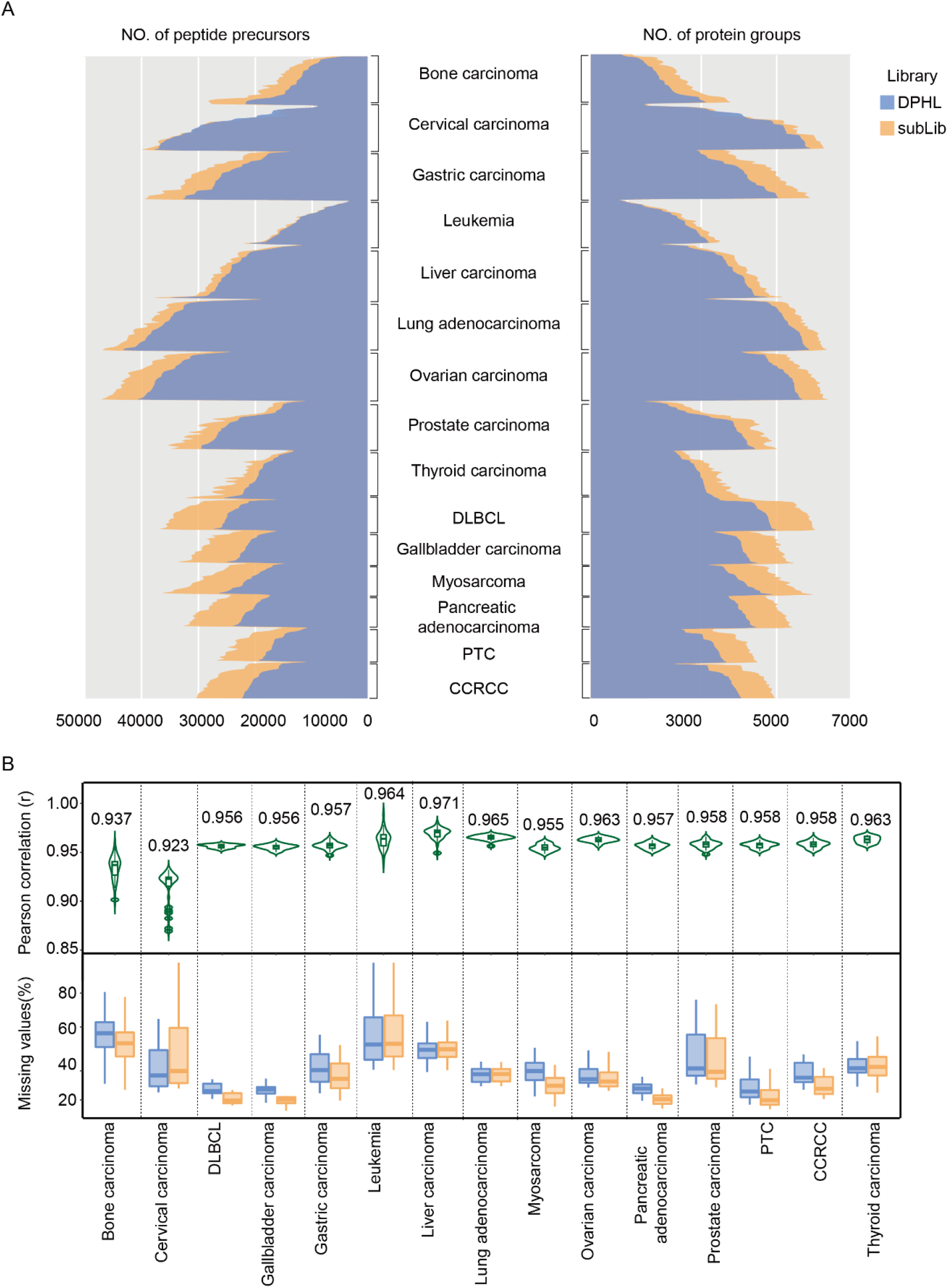
Peptide precursor and protein identification using the optimized subset library and DPHL. (A) The number of peptide precursors and protein groups identified using the optimized subset library and DPHL for each sample of every tumor type. subLib, the optimized subset library. Protein identifications were shown on the right, and peptide precursor identifications were shown on the left. (B) The correlation values on the protein level between identification results of the optimized subset library and DPHL. The percentages of protein missing values identified base on DPHL and the optimized library of each tumor type.

### Concluding remarks

In this study, we present a computational strategy to optimize library size for DIA data analysis. In our DIA data of human tissue specimens, setting FDR to 0.05 enabled effective spectral library subsetting. The application of this strategy to DIA data from 15 tumor types further consoidated this conclusion. This subLib strategy reduced false positive identifications, increased peptide and protein identifications, and generated protein data matrix quantitatively comparable to the DIA analysis with unfiltered library. In conclusion, the subLib strategy for DIA spectral library size optimization boosts proteome identifications of DIA-MS data.

## Acknowledgements

This work is supported by grants from National Key R&D Program of China (No. 2020YFE0202200), the National Natural Science Foundation of China (81972492, 21904107), Zhejiang Provincial Natural Science Foundation for Distinguished Young Scholars (LR19C050001), Hangzhou Agriculture and Society Advancement Program (20190101A04), and Westlake Education Foundation. We thank Westlake University Supercomputer Center for assistance in data storage and computation.

## Conflict of interest statement

The research group of Tiannan Guo is partly supported by Pressure Biosciences Inc, which provided access to advanced sample preparation instrumentation. T.G is shareholder of Westlake Omics Inc. W.G. is employee of Westlake Omics Inc. The remaining authors declare no competing interests.

## Data Availability

The raw data and peptide/protein matrixes were deposited in ProteomeXchange Consortium (https://www.iprox.org/). Project ID: IPX0002439000 and IPX0001981000.

---

**Figure S1.**
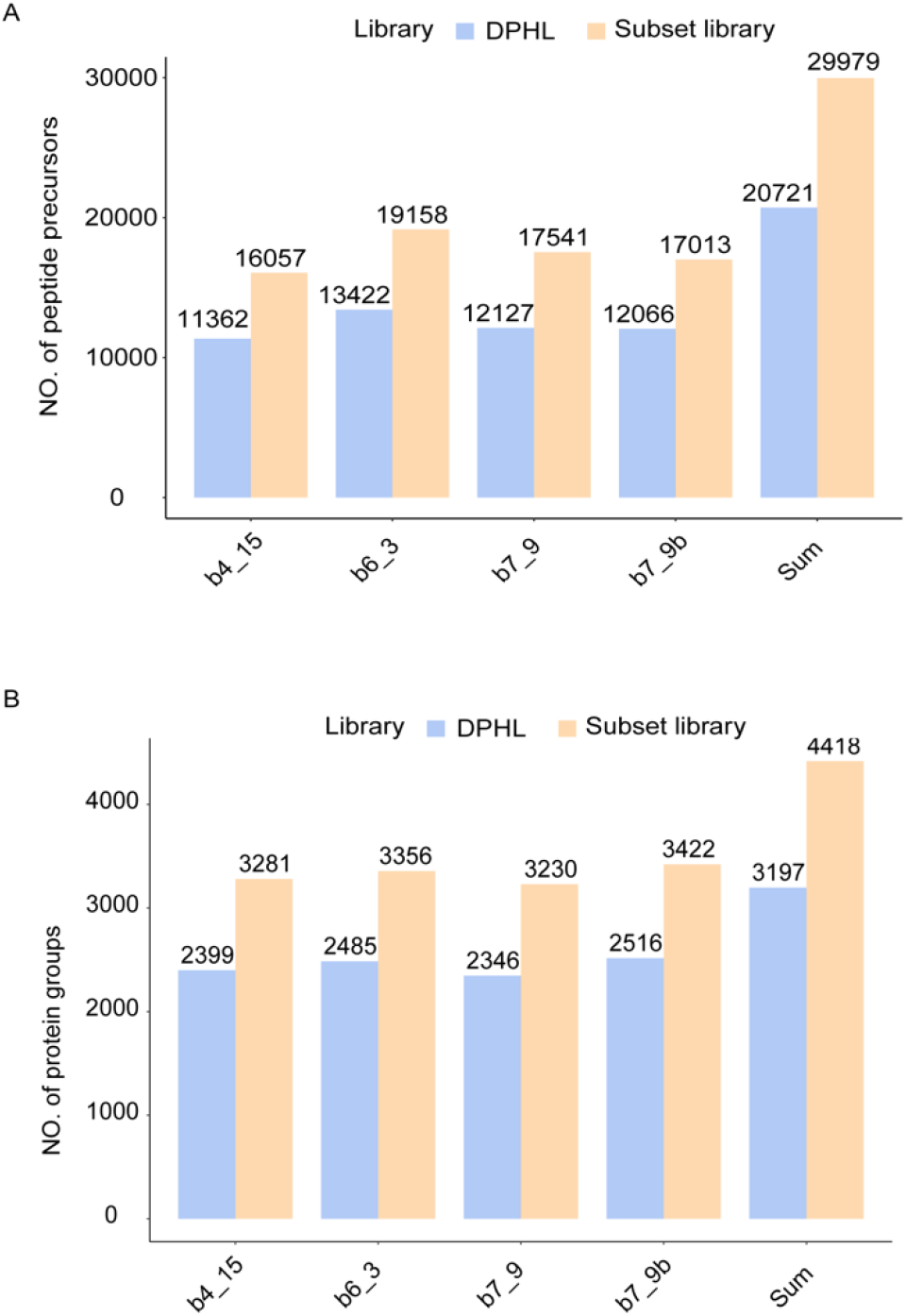
Identification results of the four representative DIA-MS data in the colorectal cancer cohort.

**Figure S2.**
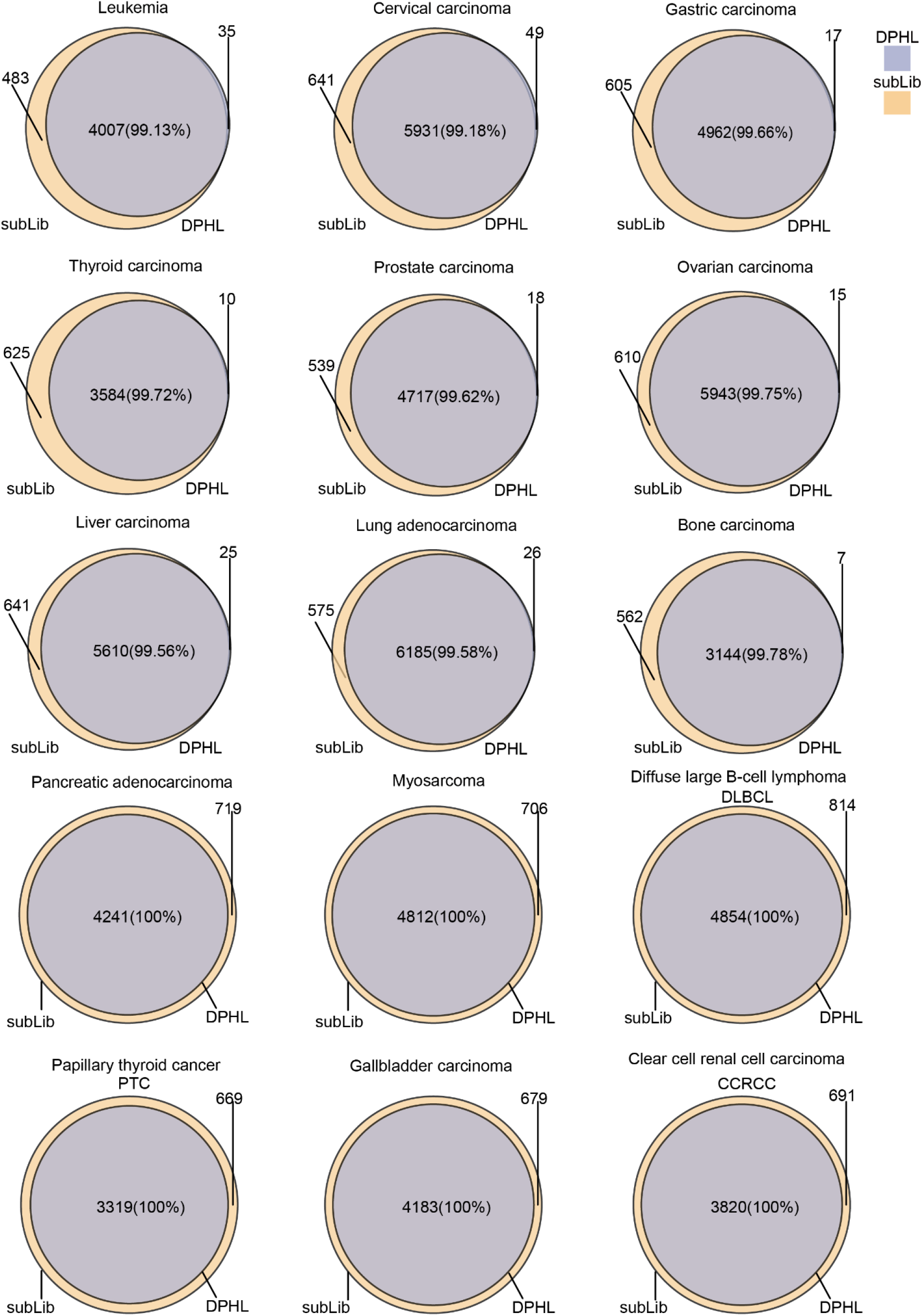
Venn diagrams showing overlap of protein identifications between the optimized subset library and DPHL.

## Notes

### Competing Interest Statement

The authors have declared no competing interest.

